# MetaNetX/MNXref - unified namespace for metabolites and biochemical reactions in the context of metabolic models

**DOI:** 10.1101/2020.09.15.297507

**Authors:** Sébastien Moretti, Van Du T. Tran, Florence Mehl, Mark Ibberson, Marco Pagni

## Abstract

MetaNetX/MNXref is a reconciliation of metabolites and biochemical reactions providing cross-links between major public biochemistry and Genome-Scale Metabolic Network (GSMN) databases. The new release brings several improvements with respect to the quality of the reconciliation, with particular attention dedicated to preserving the intrinsic properties of GSMN models. The MetaNetX website (https://www.metanetx.org/) provides access to the full database and online services. A major improvement is for mapping of user-provided GSMNs to MXNref, which now provides diagnostic messages about model content. In addition to the website and flat files, the resource can now be accessed through a SPARQL endpoint (https://rdf.metanetx.org).

## INTRODUCTION

MetaNetX/MNXref provides computed cross-references between metabolites as well as biochemical reactions, to reconcile major public biochemical databases and a selection of public genome-scale metabolic networks (GSMN) from a few dedicated resources (1–12). The previous releases of the MetaNetX/MNXref resource and website have already been presented (13, 14) as well as the main challenges of the reconciliation (15, 16, 17). This paper focuses on the current status of the resource and the recent improvements accomplished through the complete redesign and rewriting of its production pipeline.

GSMN reconstruction is one of the foundations of Systems Biology. The development of such reconstruction involves the integration of knowledge about reactions and metabolites from the scientific literature, public databases and previously published GSMNs. Historically, many GSMNs were formulated using metabolites represented as ‘symbols’, *i.e.* without explicit reference to a molecular structure. The initial motivation for creating this resource ten years ago was to add molecular structures to existing GSMNs. More generally, the goal was to establish cross-links between symbols in GSMNs published by different groups and the molecules found in the major biochemical databases. This problem is trivial as long as one-to-one mappings can be established between the different resources, but this is not always the case. Merging metabolites in a metabolic network may lead to the merging of reactions, possibly altering the model properties, and hence the predictions that could be made using it.

## MATERIAL AND METHODS

The computation of the reconciliation was performed using various different software components glued together by Bash, Perl 5.24.0 (18) and R 3.6.1 (19) scripts. These components include InChI 1.0.5 (20), Chemaxon MarvinBeans 19.26 (21), Gurobi 9.0.2 (22), Blast 2.7.1+ (23), MATLAB (24), COBRA Toolbox 2.0.0 (25) and LibSBML 5.1.8.0 (26). The MetaNetX website backend is written in Perl with the help of Apache (27) and Virtuoso (28). The frontend uses jQuery (29) and DataTables (30) with some of their plugins to ease browsing.

## RESULTS

### Principles

For MetaNetX/MNXref, a metabolite identifier (*e.g.* **MNXM123**) first refers to a *set* of metabolites in external databases, which are merged together because they are assumed to be the same biochemical entity, *i.e.* distinctions in protonation states, tautomeric forms or isotopes are typically ignored. Secondly, for every set of metabolites, a single external database identifier is selected as the (best) reference, from which a molecular structure is possibly retrieved. ChEBI identifiers that appear in Rhea reactions are preferentially selected as reference.

A consequence of the merging of metabolites is the merging of reactions involving these metabolites (Fig. 1). A reaction identifier (*e.g.* **MNXR456**) designates a *set* of reactions in external databases that are assumed to be the same biochemical entity, because they were merged together. A single external identifier is selected as the reference for this set, preferably a reaction from Rhea when feasible. However, a reaction rewritten with MNXref identifiers may differ from the original equation, for example, with respect to proton balancing. In addition, since release 2.0, a distinction was introduced between transported protons that account for the proton motive force (**MNXM01**) and those introduced to balance chemical equations (**MNXM1**). Such a distinction is also made in some recently published models (31).

**Figure 1.**
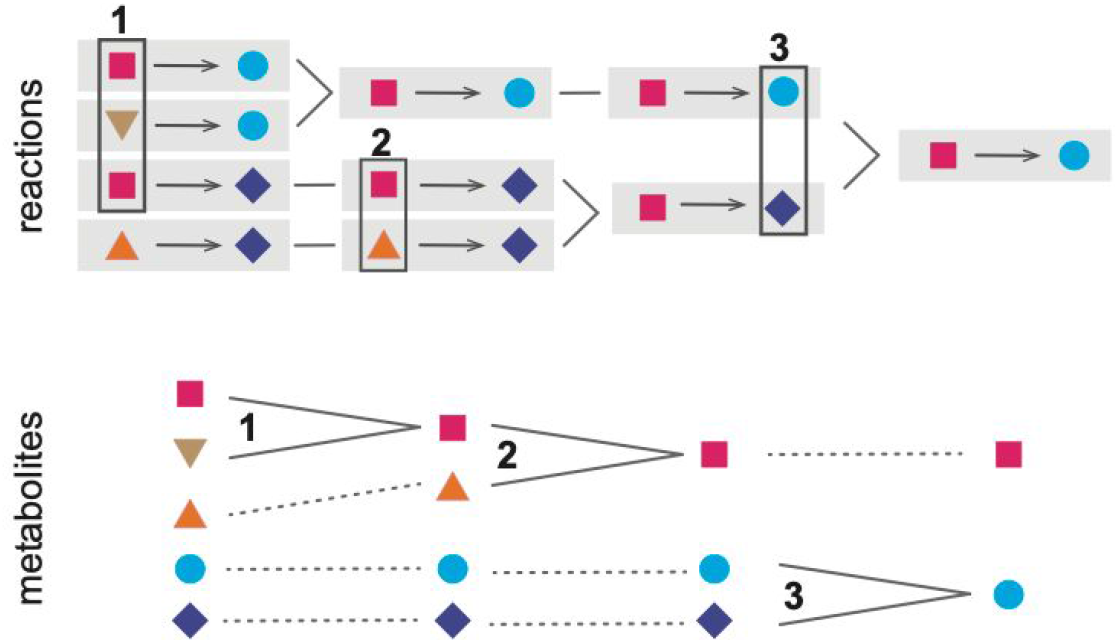
Example of the progressive reconciliation of five metabolites in four reactions into two metabolites in a single reaction. Three lines of evidence are used successively: (1) identical structures; (2) similar structures in reaction context; (3) structure and symbol pair in reaction context.

### Reconciling metabolite and reactions

Molecular structures are systematically exploited to reconcile metabolites. The proportion of chemicals in external databases endowed with a molecular structure have steadily increased over the last ten years. However, many difficulties remain, such as incomplete information on cis/trans- and stereo-isomerisms. The interpretation of partially defined structures, especially those involving R-groups and polymers, also remains a challenge.

The availability of cross-references between reactions often enable the reconciliation of pairs of metabolites if supported by additional evidence, for example the compound name. This reconciliation of metabolites from the *reaction context* evidence was the key idea that allowed the first computation of MetaNetX/MNXref (15). It is to be noted that imported cross-references between metabolites are ignored during the reconciliation, to avoid propagating existing errors present in external databases.

### Manual curation

Although we aimed to make the reconciliation procedure fully automated, errors are occasionally imported from external resources that can affect the reconciliation procedure. Automated detection of conflicting evidence and ambiguities has been central to our effort to rewrite the pipeline. Thereafter, manual case-by-case study enables fixing of flagged issues, for example by correcting the reference of a metabolite to a molecular structure. An archetypal example of such difficulties are the two stereo-isomers of a compound - according to the textual description - where the structure of the L-form is fully specified and the structure of the R-form misses the stereochemistry.

Two metabolites to be merged that appear in the same reaction, in the left and right terms for example, also deserve scrutiny. Pairs of tautomers or acid-bases can be validly merged, possibly yielding an *empty* reaction that will disappear from the mapped GSMN. Changes in reaction status (see below) has also been used as a trigger for the manual inspection of a particular reconciliation.

The errors we have detected have been systematically reported to the resource from which they originated. Most of them have since been corrected, thanks to the diligence of their curators.

### Impact of reconciliation on model properties

One of the main challenges of the MNXref reconciliation is to ease comparison of GSMNs while preserving their intrinsic *properties.* We have attempted to summarize the latter by attributing a *status* to every reaction in a GSMN (Fig. 2): **A**, the reaction can carry a non-zero flux in the unaltered GSMN; **a,** the reaction can carry a flux after all boundary (exchange) reactions were set to bi-directional; **b,** the reaction can carry a flux after all reactions in the GSMN were set to bi-directional; **B**, the reaction cannot carry a flux because of the network topology, for example because of a dead-end metabolite. Computation is performed using a variant of the flux variability algorithm (32) with limited numerical precision for GSMN with more than 5000 reactions.

**Figure 2.**
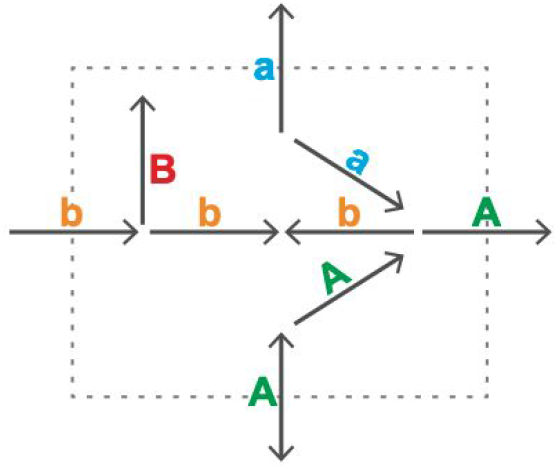
A status (A, a, b, B) was attributed to every reaction of this simple network, as explained in the text. A reaction status ultimately depends on the whole network topology and the directionality of every reaction.

For example, the iAF1260 model for *E. coli* (*33*) contains a total of 2376 reactions. After mapping to MNXref, 2368 reactions are retrieved with a one-to-one mapping to an original reaction. The status of these reactions is distributed as follows: **A**, 1518; **a**, 627, **b**, 87; **B**, 136. The reaction status is perfectly conserved in this example, despite the merging of two metabolites namely **1agpg140** (1-Acyl-sn-glycero-3-phosphoglycerol (n-C14:0)) and **1tdecg3p** (1-tetradecanoyl-sn-glycerol 3-phosphate). In other GSMNs, the merging of metabolites may cause the status of a few reactions to change, the latter not necessarily involving the merged metabolites and possibly augmenting the number of reactions that can carry a non-zero flux. Taking control of such indirect effects has been crucial for testing and improving the reconciliation. More public GSMNs will be added in the future to further challenge and improve the quality of the reconciliation.

### Database content

Tables 1 and 2 list the different resources that were included in the latest release. The ratio of the numbers of entries before and after reconciliation is a measure of its efficiency. Indeed, this ratio is most optimal for the metabolites and reactions found in around hundred GSMNs hosted at MetaNetX.

**Table 1.**
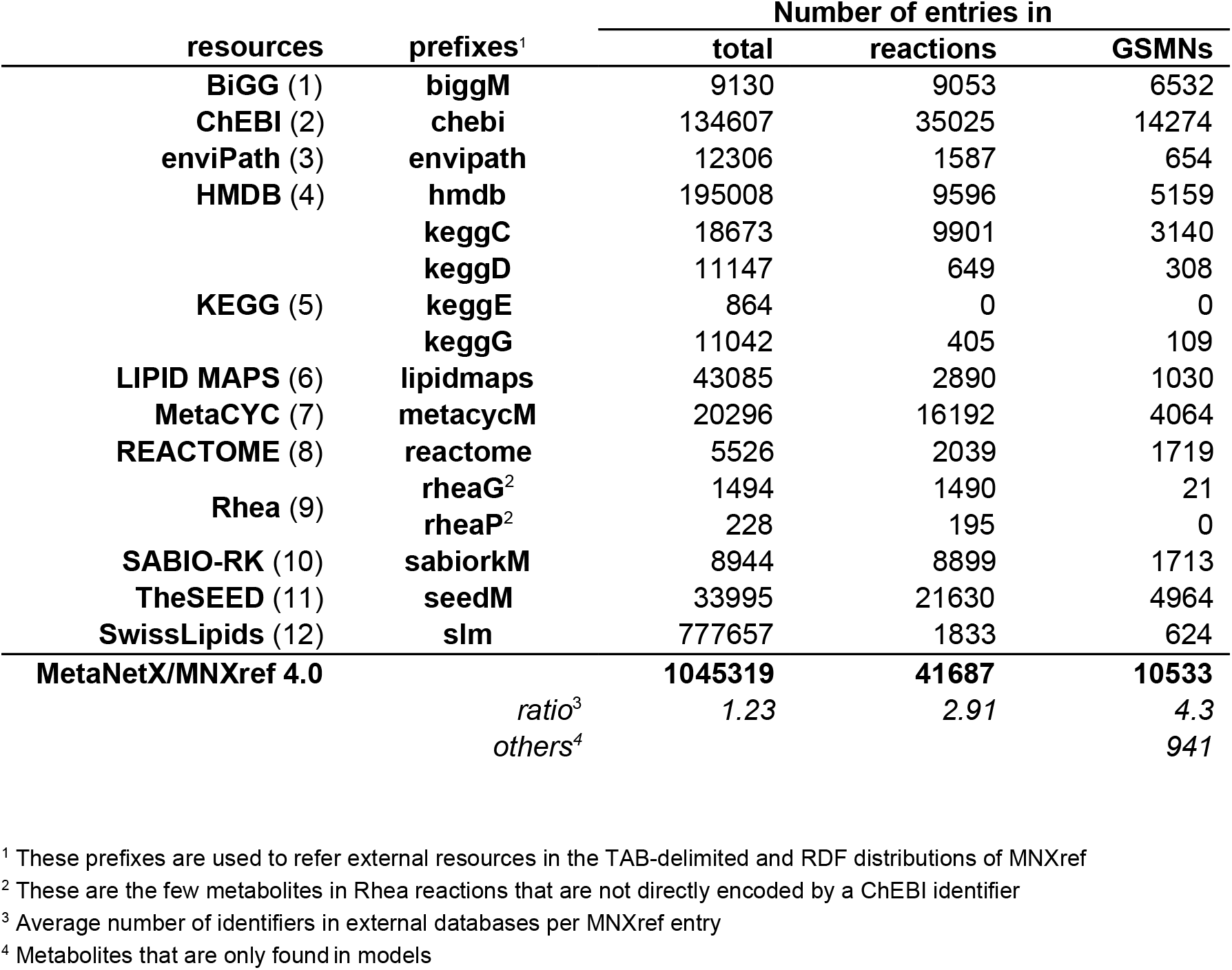
Sources and numbers of metabolites found in the MNXref 4.0 reconciliation. The number of metabolites found in the different resources and how non-redundant they are made through MNXref reconciliation. Secondary and deprecated identifiers are not counted here.

**Table 2.** Sources and numbers of reactions found in the MNXref 4.0 reconciliation. Secondary and deprecated identifiers are not counted here.

| resources | prefixes<sup>1</sup> | Number of entries in |  |
| --- | --- | --- | --- |
|  |  | total | GSMs |
| **BiGG** (1) | **biggR** | *28167 | *41863 |
| **KEGG** (5) | **keggR** | 11161 | 3508 |
| **MetaCYC** (7) | **metacycR** | 17208 | 4898 |
| **Rhea** (9) | **rheaR**<sup>2</sup> | 12510 | 3989 |
| **SABIO-RK** (10) | **sabiorkR** | 8118 | 2319 |
| **TheSEED** (11) | **seedR** | *43855 | *19230 |
| **MetaNetX/MNXref 4.0** |  | **37103** | **11968** |
|  | *ratio*<sup>3</sup> | 3.26 | 6.87 |
|  | others<sup>4</sup> |  | 6463* |

\* Reactions with specific compartments (vs generic compartments)
<sup>1</sup> These prefixes are used to refer external resources in the TAB-delimited and RDF distributions of MNXref
<sup>2</sup> Only master reactions of Rhea are counted here
<sup>3</sup> Average number of identifiers in external databases per MNXref entry
<sup>4</sup> Reactions only found in models, but most of them are formulated with metabolites in MNXref

### Service

The MetaNetX website (https://www.metanetx.org) provides access to the full database and online services. Most notably, the service that permits a user to upload his or her own GMSN and map it to MNXref has been greatly improved to produce diagnostic messages about the model content. The database can be queried through the website user interface, as well as through a SPARQL endpoint (https://rdf.metanetx.org).

## DISCUSSION

By design a GSMN is a model of the metabolism of low molecular weight compounds, where mass conservation and thermodynamics apply (34, 35). GSMN models play a crucial role in the toolbox of synthetic biology (36, 37). Beyond popular applications, such as metabolic flux prediction with flux balance analysis and prediction of the gene essentiality, GSMNs have been used in numerous applications (38), e.g. chemicals and materials production (39–42), drug targeting (43, 44), human metabolism, disease understanding (45, 46) and, multi-organism interaction modeling (47, 48). MNXref is now used within tools for testing GSMNs such as MEMOTE (49), and has been proposed as a reference for minimal standard content for metabolic network reconstruction (50). Finally, the ability to accurately reconcile metabolites and reactions within GSMNs paves the way for applications such as multi-omics data interpretation (51, 52), and white-box AI models (53). Moving forward, MetaNetX/MNXref efforts to not only unify metabolites, reactions and subcellular compartments but also genes and proteins (54) into a single comprehensive namespace will continue to provide a strong basis for such key systems biology endeavors in the future.

## AVAILABILITY

Data is distributed under a Creative Commons Attribution 4.0 International License (**CC BY 4.0**). The MNXref reconciliation is available in TAB-delimited and RDF/Turtle formats. The mapped GSMNs are available in SBML level 2 and 3 formats from the web site and as part of the RDF graph.

## ACKNOWLEDGEMENT

Computation and maintenance of the MetaNetX.org server are provided by the Vital-IT (https://www.vital-it.ch) and Core-IT centers of the SIB Swiss Institute of Bioinformatics (https://sib.swiss). We thank Alan Bridge, Anne Morgat and the whole Rhea team for stimulating exchanges and discussions. We thank Jerven Bolleman for convincing us that RDF was an effective technology. We thank Manuela Pruess for help with the manuscript.

